# Enriching human interactome with functional mutations to detect high-impact network modules underlying complex diseases

**DOI:** 10.1101/786798

**Authors:** Hongzhu Cui, Suhas Srinivasan, Dmitry Korkin

## Abstract

Progress in high-throughput -omics technologies moves us one step closer to the datacalypse in life sciences. In spite of the already generated volumes of data, our knowledge of the molecular mechanisms underlying complex genetic diseases remains limited. Increasing evidence shows that biological networks are essential, albeit not sufficient, for the better understanding of these mechanisms. The identification of disease-specific functional modules in the human interactome can provide a more focused insight into the mechanistic nature of the disease. However, carving a disease network module from the whole interactome is a difficult task. In this paper, we propose a computational framework, DIMSUM, which enables the integration of genome-wide association studies (GWAS), functional effects of mutations, and protein-protein interaction (PPI) network to improve disease module detection. Specifically, our approach incorporates and propagates the functional impact of non-synonymous single nucleotide polymorphisms (nsSNPs) on PPIs to implicate the genes that are most likely influenced by the disruptive mutations, and to identify the module with the greatest impact. Comparison against state-of-the-art seed-based module detection methods shows that our approach could yield modules that are biologically more relevant and have stronger association with the studied disease. We expect for our method to become a part of the common toolbox for disease module analysis, facilitating discovery of new disease markers.

## 1. Introduction

Since the first evidence of the genetic complexity in cancer [1], numerous research efforts have been dedicated to deciphering the mechanistic nature of polygenic diseases. Due to the rapid advancement of the next generation sequencing (NGS) technologies, including whole-genome [2] and whole-exome sequencing [3], and most recently single-cell transcriptomics [4], we have been able to sequence and analyze thousands of genomes at a much lower cost. As result, the high-throughput experiments produce large amounts of genomic data at the exponentially increasing rate. These data have also transformed the design of genome-wide association studies and enabled us to do comprehensive analyses of the genotype-phenotype relationships [5]. Most importantly, the studies have provided us with an extensive list of susceptible alleles and genes associated with complex diseases as well as with catalogs of disease-relevant mutations[6]. The list includes the majority of common and many rare complex diseases. And it is expected that a comprehensive catalog of nearly all human genomic variations will be available soon [7].

A typical large-scale NGS-based study reports several million genetic variants [5]. However, not all mutations, even with statistically significant correlations, would contribute to the disease phenotype [8]. For example, in cancer genomics, many mutations are so-called passenger mutations. Unlike the driver mutations, which induce the clonal expansion, the passenger mutations do not provide any functional advantage to the development of cancer cells [9, 10]. Thus, distinguishing between the functional and non-functional mutations is usually the first step in genetics studies. Additionally, computational approaches for functional annotation are increasingly important, since many variants are not previously described in the literature [6, 11]. There are a plethora of functional annotation tools for genetic variants [6]. Many tools focus on the annotation of single nucleotide variants (SNVs), the most common type of genetic variation, which is easier to capture and analyze. Recently, we have developed a new computational method, SNP-IN tool [12], which predicts the effects of non-synonymous SNVs on protein-protein interaction (PPI), provided the interaction’s structure or structural model. The method leverages supervised and semi-supervised feature-based classifiers, including a new random forest self-learning protocol. The accurate and balanced performance of SNP-IN tool makes it apt for functional annotation of non-synonymous single nucleotide polymorphisms (nsSNPs). SNP-IN tool could also be helpful system-wide variation analysis to discover pathways shared by associated alleles and reveal disease-related biological process.

The need of integrating GWAS studies and the functional impact of the disease-associated mutations with the systems data is supported by the increasing body of evidence that large-scale biological systems and cellular networks underlie majority—if not all—of the complex genotype-phenotype relationships in diseases [13-16]. Understanding the biological network is essential in studying a genetic disease, because such a disease is likely a result of the disruption and rewiring of the complex intracellular network, rather than a consequence of the dysfunction of a single gene [17]. Along with the increasing availability of high-throughput human protein interactomics data [18, 19], computational approaches have been developed. In particular, network propagation has recently emerged as a prominent approach in network biology [20]. In network propagation, genes/proteins of interest correspond to the nodes in the biological network, the edges represent pair-wise protein-protein interactions, and the information is propagated through the edges to nearby nodes in an iterative fashion. Thus, it could amplify the weaker disease association signals from the genes interacting with the “seed” genes that carry the stronger source signal [20]. Network propagation have been applied for various purposes, including predicting gene function, identifying disease related subnetworks, and drug target prediction [20, 21].

Module identification is a central problem in network biology [22]. A topological module is a set of genes (nodes) with dense interactions between each other; these groups of nodes are also referred to as communities, or clusters, in network science [23]. A topological module can be functional, since the constituting proteins are often shown to pertain to the same biological function or be involved in a similar biological process. A disease module is a sub-network of proteins enriched with the disease-relevant proteins and responsible for the disease phenotype. Various module identification approaches have been proposed, presenting a wide range of theoretical perspectives and implementations. These methods primarily come in two different flavors. The first group of methods identify the modules in a biological network by relying exclusively on the network’s topology. This is a challenging task due to the lack of information about specific genes/proteins contributing to biological functionality or disease phenotype. The recent open community DREAM challenge [24] provided a good review and benchmark of existing methods falling in this category. Methods from the second group start with the “seed genes”, and gradually extract additional genes in the network to grow the module. For example, DIAMOnD [25] is a disease module detection algorithm that utilizes known seed genes to identify disease modules according to the number of connections to the seed proteins. The algorithm outputs a connected disease module with a list of candidate disease-associated proteins ranked by their connectivity significance.

Discovering biologically relevant modules (disease modules or functional modules) is a challenging task [26, 27]. In fact, to tackle the mechanistic intricacy underlying complex diseases, it is necessary to couple disparate sources of data, each informing about a different aspect of the biological function. Computational approaches integrating molecular networks with different types of -omics data have demonstrated considerable power in bioinformatics studies [28]. Integrating PPI networks with gene expression profiles to identify sets of genes that participate in a biological function is a successful example of the benefits of data integration [29]. Interestingly, integrating interactomics data with GWAS data does not garner wide attention in the bioinformatics community, and functional annotation information regarding genetic variants and mutated genes are mostly missed out during such integration process.

In this work, we developed a novel algorithmic framework named DIMSUM (<u>D</u>iscovering most <u>IM</u>pacted <u>S</u>Ubnetworks in interacto<u>M</u>e) to identify functional disease module in the human interactome. The DIMSUM framework includes three major steps: 1) network annotation; 2) network propagation, and 3) subnetwork extraction. Our approach benefits from integrating GWAS data, functional annotation information, and the protein-protein interaction network. We evaluated our approach using a set of 8 complex diseases against two state-of-the-art seed-based module detection methods: DIAMOnD and SCA. From the set of complex diseases, we also carried out two case studies: the first study centered around genes associated with coronary artery disease, and the second one focusing on joint analysis of Schizophrenia and Bipolar Disorder. The evaluations results show that DIMSUM outperforms both DIAMOnD and SCA, because the discovered modules have stronger association with the disease and are more biologically relevant with the seed-gene pool.

## 2. Materials and Methods

At the preprocessing stage, we carry out the human interactome construction, GWAS data collection, and data processing (Fig. 1). At the same time, we map the mutations to the structures of affected PPIs, and apply our SNP-IN tool[12] to characterize mutation-induced rewiring effects on the PPIs[13]. The computational workflow of DIMSUM consists of 3 main stages. In the first stage, one updates the node and edge weights of a fully annotated human PPI network: the node weight reflects the association with a disease, while the edge weight reflects the cumulative damage made to the corresponding interaction. Second, we apply a network propagation strategy with a goal to boost the signal for the genes from GWAS study with weak association, thus increasing the pool of candidate genes. The last stage includes sub-network extraction: we propose an iterative procedure to find the disease module with the greatest impact on the disease.

**Figure 1.**
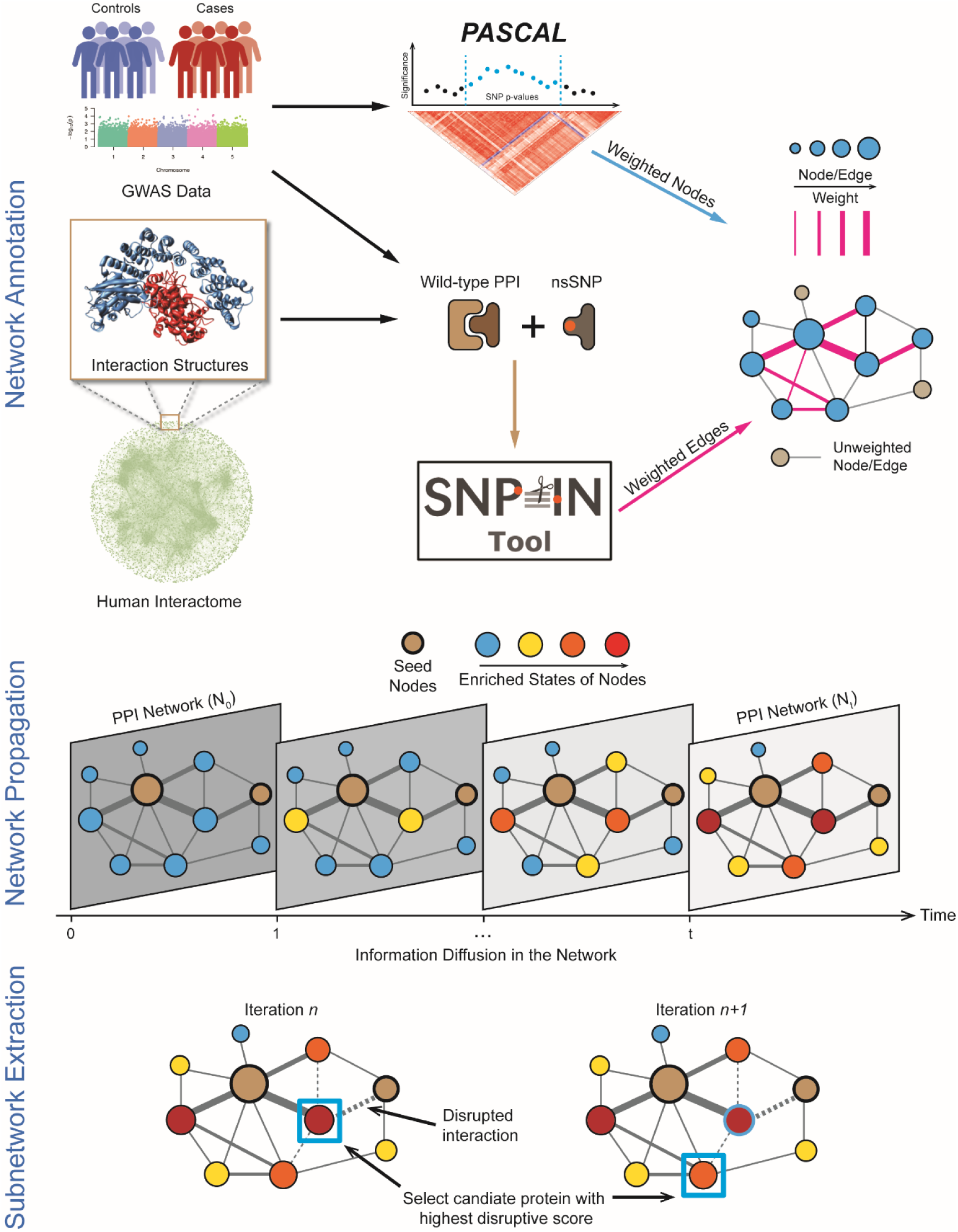
Overview of the workflow of DIMSUM. The module detection framework contains three major steps: network annotation; network propagation & subnetworks extraction. Network annotation: collect GWAS data and use Pascal tool to aggregate SNP summary statistics; apply SNP-IN tool to properly assess SNP’s impact on the protein-protein interactions. Integrate this information with the human interactome to generate a fully weighted network. **N**etwork propagation: apply a network propagation procedure to implicate genes most likely to be influenced by disruptive SNPs. Subnetwork extraction: apply an iterative strategy to identify the most impacted module.

### 2.1. Human interactome construction

In this work, we utilize two different protein-protein interaction data sources to construct the human interactome. The first data source is the HINT database [30]. It integrates several databases and filters out low-quality and erroneous interactions. The other source is the Human Reference Protein Interactome Mapping Project (HuRI) [19]. HINT is a manually curated repository of PPIs mainly from literature, whereas HuRI is a primary source of experimentally validated PPIs. We consider these two PPI sources as they complement each other, hence we construct the interactome by combining both. The current release of HINT interactome (Version 4) contains 63,684 interactions. Combining all the three proteome-scale human PPI datasets released from HuRI at different stages of the project, we obtain 76,537 interactions. In total, we generate a human interactome consisting of 105,087 interactions, where 35,134 interactions exist in both data sources.

### 2.2. GWAS data collection and processing

A distinctive feature of our method is that it integrates GWAS data into the interactome to improve the disease module detection. To do so, we compiled eight publicly available GWAS datasets. The datasets span a broad range of diseases, including neurodegenerative, metabolic, and psychiatric disorders. For the GWAS integration, the dataset is only required to have pre-calculated summary statistics, no individual level information is needed.

To generate the seeds for the later stage of network propagation, we compute gene scores by aggregating SNP *p*-values from GWAS studies using the *Pascal* tool (<u>Pa</u>thway <u>sc</u>oring <u>al</u>gorithm) [31]. Integrating SNP *p*-values from GWAS studies has proven itself as a powerful method to improve statistical power.

Pascal is a fast and rigorous computational tool developed to aggregate SNP summary statistics into gene scores with the high power, while absolving the need to access the original, individual-level, genotypic data. It can be considered as an alternative to the traditional *p*-value estimation approaches, including chi-squared statistics (SOCS) and the maximum of chi-squared statistics (MOCS) [32], which measure the average and the strongest associations of signals per gene, respectively. Pascal relies on the assumption that a pairwise correlation matrix of the contributing genotypes underlying the null distributions of the MOCS and SOCS statistics can be estimated from the ethnicity-matched, publicly available genotypic data. To calculate gene scores, we select the sum of chi-squared statistics (SOCS) of all SNPs from genes of interest. To properly correct for linkage disequilibrium (LD) correlation structure in GWAS data, we use the European population of the 1,000 Genomes Project [33], because GWAS studies in this work are predominantly the European cohorts. The disease-associated genes with significant p-values that Pascal produces are then defined as the seed genes for our approach. The final list of seed genes is acquired after correcting for multiple testing using Bonferroni correction.

### 2.3. Functional annotation of nsSNV with SNP-IN tool

To properly assess the functional damage caused by mutations with respect to protein-protein interactions, we apply our recently developed SNP-IN tool (non-synonymous SNP INteraction effect predictor tool) [12]. SNP-IN tool predicts the rewiring effects of nsSNVs on PPIs, given the interaction’s experimental structure or accurate comparative model. More specifically, SNP-IN tool formulates this task as a classification problem. There are three classes of edgetic effects predicted by the SNP-IN tool: beneficial, neutral, and detrimental. The effects are assigned based on the difference between the binding free energies of the mutant and wild-type complexes (ΔΔG). Specifically, ΔΔG = ΔΔG_*mt*_ - ΔΔG_*wt*_, where ΔΔG_*mt*_ and ΔΔG_*wt*_ are the mutant and wild-type binding free energies correspondingly. The beneficial, neutral, or detrimental types of mutations are then determined by applying two previously established thresholds to ΔΔG [34, 35]:

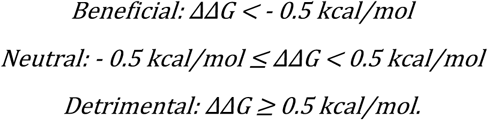

The annotation workflow begins with processing the GWAS data. Most mutations in GWAS datasets come with only dbSNP RefSNP cluster ID’s (rs#). The variant data is preprocessed using ANNOVAR [36] to retrieve SNV locations on the genes and the corresponding residue change information. For each mutated gene and the corresponding SNP, we collect all their interacting partners in the merged interactome (see Section 2.1). Because SNP-IN tool is a structure-based classifier, we need both the mutation information and interaction structure as an input. There are several different cases when generating the PPI structure, in which a mutated gene is involved (Supplementary Fig. 1). First, if a PPI already has a native structure, it is extracted from protein data bank (PDB) [37]. In the corresponding PDB file, we first identify the interacting subunit pair for each PPI using 3did database [38]. The 3did database maintains information regarding the two interacting domains with physical interfaces. If there is no native structure for a protein-protein interaction, two options are explored. First, if a structural template for such interaction (*i.e*., a homologous protein interaction complex) exists, a comparative model of this interaction can be obtained [39]. When a full-length PPI cannot be modeled, we only model the domain-domain interaction that includes the domain containing the mutation. Homology modeling is done through Interactome3D [40], a web service for structural modeling of PPI network.

### 2.4. Network annotation and network propagation

The human interactome is next represented as the graph *G* = (*V, E*), where *V* is the set of nodes representing the genes; *E* is the set of edges representing the protein-protein interactions. After applying the Pascal tool to the GWAS studies, we obtain *p*-values of disease associated genes, these values together with the functional annotations of the corresponding mutations from the SNP-IN tool, are used to weight the nodes and edges in the network. Thus, *G* is a graph with weighted nodes and edges. Specifically, for a disease-associated gene *i*, the corresponding node is weighted with –log(*P*_*i*_), where *P*_*i*_ is the *p*-value obtained from Pascal tool. A node corresponding to a gene that is not listed in the GWAS study is assigned a zero weight. The edge is weighted according to the damage accumulated on the corresponding PPI by the disruptive SNVs. Specifically, a PPI between genes *i* and *j* is weighted with the total number of disruptive SNVs on both genes targeting the same interaction. In other words, the node weight reflects the “relevance” of the gene to the disease, and the edge weight reflects the accumulated “damage” on the interaction.

Next, we apply the network propagation strategy to implicate other core disease genes affected by the perturbations of disruptive SNVs. Let *F: V* → ℜ represent a function reflecting the relevance of gene *i* to a specific complex disease. The goal of the network propagation is to prioritize the genes that are not showing significant association based on the GWAS study, but are expected to have possible relevance to the disease. We impose two constraints on the prioritization function *F*: the computed function should be (1) smooth and (2) comply with the prior knowledge. Smoothness of the function is defined by the assignment of similar values to the interaction partners (nodes) of disease gene *i*. The compliance with the prior knowledge implies that the difference between the final computed value for a disease gene *F(i)* and the initial value. The values of *F* can be iteratively obtained based as follows:

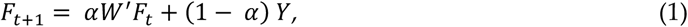

where *Y* is the prior knowledge defined as the node weight, α is a parameter reflecting the importance of the two constraints mentioned above with the default value α=0.5, *W’* is a |*V*|*|*V*| matrix whose values are determined by the edge weights and that is defined as a normalized form of network edge weight matrix *W*. Formally, we introduce a diagonal matrix *D*, such that *D(i, i)* is the sum of row *i* of W. We then set *W’ = D*^*-1/2*^*WD*^*-1/2*^. *F*_*t*_ is initialized as *Y*. The equation could be solved iteratively and guaranteed to converge to the system’s solution [41]. This iterative algorithm can be considered as propagating the prior information from some nodes through the network. The disease genes first send the signal to their neighbors, and every node then propagates the received signal to its neighbors (Fig. 1). There are also additional constraints on the weighted network. First, when we propagate the information through the network, both the node and edge weights should be in [0, 1] range [41]. In this work, the seed genes in the network carry the largest weights. Thus, we normalize the node weight:

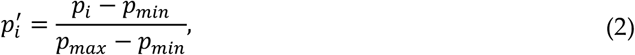

where *p*_*max*_ and *p*_*min*_ are the maximum and minimum of the initial node weights, *i.e*. –log(*P*_*i*_). After network propagation, we de-normalize the node weight at the subnetwork extraction step (see Section 2.5 next). For the edge weight, the higher weight means that the information flow is more likely to go through that edge. Thus, the weight will be converted to [0,1] range using a sigmoid transformation:

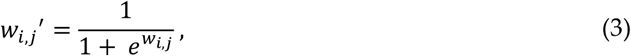

where the *w*_*i,j*_ is the original weight of the edge. After convergence, the disease association information from seed genes is diffused into the interactome, and all the nodes are weighted with their relevance to the disease.

### 2.5. Sub-network extraction

Finally, we extract the sub-network with the greatest “impact” on the disease etiology. To do so we define disease-associated genes with significant *p*-values obtained from Pascal as the seed genes. The goal is to extract a sub-network containing all the seed genes, while maintaining the greatest impact. Intuitively, the impact is defined based on the “severity” of the network damage caused by the disruptive SNVs located on the genes with high relevance to the disease. The relevance is reflected by the node weight after the network propagation procedure, while the network damage is determined based on the edge weights reflecting the total number of disruptive mutations occurring in each interaction. The subnetwork extraction is then formulated as an iterative procedure:

I. Assume that a seed gene set {*g*_1_, *g*_2_, …, *g*_*k*_} induces a subnetwork with initial size *N*_0_. For all immediate interacting partners of the seed gene set that are not in the subnetwork, define an impact score:

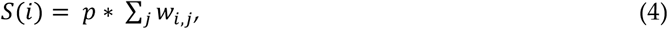

where *p* is the updated *p*-value after the network propagation and ∑_*j*_ *w*_*i,j*_ is the total number of mutations that disrupt the PPIs between the candidate gene *i* and every gene *j* in the module gene set. Thus, the impact score combines the disease relevance and potential disruption to existing subnetwork caused by the candidate gene.
II. All the immediate interaction neighbors of the seed gene set, not included in the subnetwork, are ranked according to their impact score.
III. Select the gene with the largest impact score. If there are multiple candidates with the same impact score, randomly pick one gene to break the tie. Add the gene with the biggest impact to the set of seed genes and increase the size of the induced subnetwork by 1: *N*_*t*+1_ = *N*_*t*_ + 1.

Given a number of maximum iterations, steps I - III are repeated in a loop until the number of added genes equals maximum iteration number. The code for network propagation and sub-network extraction is available at: https://github.com/hcui2/DIMSUM.

### 2.6. Validation and GO analysis

To validate the performance of our module detection method, we compared it against two seed-based module detection methods, a widely used method called DIAMOnD [40], and a recently published method SCA [42]. For both methods, we used the same seed genes that were used for DIMSUM. For each method, we also limit the number of candidate genes forming a disease module to 100. To validate the disease association of the predicted candidate genes, we first compiled a list of known disease-related genes from two databases, HGMD [42] and OMIM [43]. The candidate genes supported by the literature are considered as true positives.

Furthermore, to compare the biological relevance of candidate genes to the seed genes we perform GO enrichment analysis [44] on the two gene sets. In the GO enrichment analysis, we use the third level of the GO hierarchy as a trade-off between the too general and well-populated GO terms at the second level, and specific but not well-populated terms at the fourth level. The GO enrichment is performed using DAVID webserver [37], and GO terms with p-value ≤ 0.01 are selected. During the analysis, we first identify significantly enriched GO terms within the seed genes. We then check how many of the GO terms significantly enriched in the pool of candidate genes are identical to the enriched GO terms in the pool of seed genes. The higher number of the identical GO terms suggests the stronger relevance between the candidate genes and the seed genes.

## 3. Results

### 3.1. Seed genes generated from GWAS datasets

We collected eight GWAS datasets from various public sources. The collected GWAS data cover a diverse range of eight complex diseases, including Alzheimer’s, bipolar disorder, coronary artery disease, macular degeneration, osteoporosis, rheumatoid arthritis, schizophrenia, and type 2 diabetes mellitus (Supplementary Table 1). The GWAS datasets were only required to contain SNP-phenotype association summary statistics, no individual level genotype information was used. These studies were predominantly from European cohorts, with the total number of SNPs reported in each study varying from 2 million to 12 million (Supplementary Table 1).

**Table 1.** Comparison of the number of disease associated genes in the detected modules with literature evidence between three methods: DIAMOnD, SCA, and DIMSUM

| Disease ID | Disease name | Seeds<br>match | DIAMOnD<br>match | SCA<br>match | DIMSUM<br>match |
| --- | --- | --- | --- | --- | --- |
| D1 | Coronary artery disease | 8 | 1 | 0 | 10 |
| D2 | Diabetes mellitus - Type 2 | 14 | 0 | 1 | 2 |
| D3 | Macular degeneration | 17 | 0 | 0 | 3 |
| D4 | Osteoporosis | 5 | 1 | 1 | 0 |
| D5 | Alzheimer's disease | 8 | 0 | 0 | 3 |
| D6 | Rheumatoid arthritis | 19 | 0 | 0 | 4 |
| D7 | Bipolar disorder | 12 | 0 | 0 | 8 |
| D8 | Schizophrenia | 29 | 0 | 0 | 14 |

Our subnetwork detection strategy benefited from integrating the GWAS dataset and network data. Specifically, we derived the seed genes for network propagation from the GWAS dataset (see Subsection 2.2 in Methods). The length of the gene list generated from Pascal varied for each disease due to different sizes of GWAS datasets. The number of seeds for each disease ranged from 26 to 301 (Supplementary Table 2).

### 3.2. Functional predictions from SNP-IN tool

The lack of functional knowledge for SNPs obtained from GWAS studies limits our understanding of the mechanistic processes that underlie diseases. Although there is a plethora of functional annotation tools for SNPs, most of them provide with annotations of generic putative deleterious effects of SNPs [6]. In particular, they do not provide the means to determine how SNPs disrupt protein-protein interactions, while such information could lead to a better understanding of how SNPs rewire the human interactome and help identify the impacted subnetworks responsible for the disease. Our recently developed SNP-IN tool accurately predicts how mutations affect the PPIs, given the interaction’s structure (see Subsection 2.3 in Methods). Given that the sizes of GWAS datasets considered in this work varied, the prediction coverage of non-synonymous SNPs also varied for different diseases, ranging from 547 to 8,323 (Supplementary Table 3). Previous studies reported high percentage of disease-associated mutations that affected PPIs [13, 15, 45]. Specifically, our latest study [13] showed that out of all pathogenic mutations collected from the ClinVar database, 76.2% were predicted to have a disruptive effect on PPIs. In the present work, on average, 51.1% of the annotated mutations from all eight GWAS studies were predicted to have detrimental effects on PPIs. The percentages of disruptive mutations in this study were lower than our previous work, which could be attributed to the fact that some of mutations detected in the GWAS studies were random mutations or passenger mutations without significant functional impact. Nevertheless, the current study showed that a considerable amount of mutations occurring in a disease could rewire the human interactome centered around the disease. In addition, these results reaffirmed our previous findings that the beneficial mutations, strengthening the PPIs, were rare in the human genome (Supplementary Table 3).

### 3.3. Network annotation and network propagation

A key idea of our approach is in integrating GWAS data and functional annotation data with the human interactome data to improve the network module detection. Specifically, we utilized the GWAS study results and functional annotations from SNP-IN tool to properly weight the network. The node weight reflected the relevance of the gene to the disease, while the edge weight represented the cumulative damage imposed on the corresponding interaction (see Subsection 2.4 in Methods). Once the fully weighted network was generated, we examined the topological properties of the genes carrying disruptive mutations and the damaged interactions in the human interactome. In particular, between the eight datasets we compared the distribution of the node degrees in the human interactome for disease-associated genes. The average node degree for the genes carrying disruptive mutations among the eight diseases ranged from 14 to 24 (Fig. 2.a). Compared to the average node degree of the human interactome, all disease networks showed increased average degree suggesting that disease-associated mutations tend to disrupt genes occupying a central spot in the human interactome, rather than lying on the periphery. The results also suggested that mutations captured in complex diseases were more likely to cause the network rewiring than random mutations.

**Figure 2.**
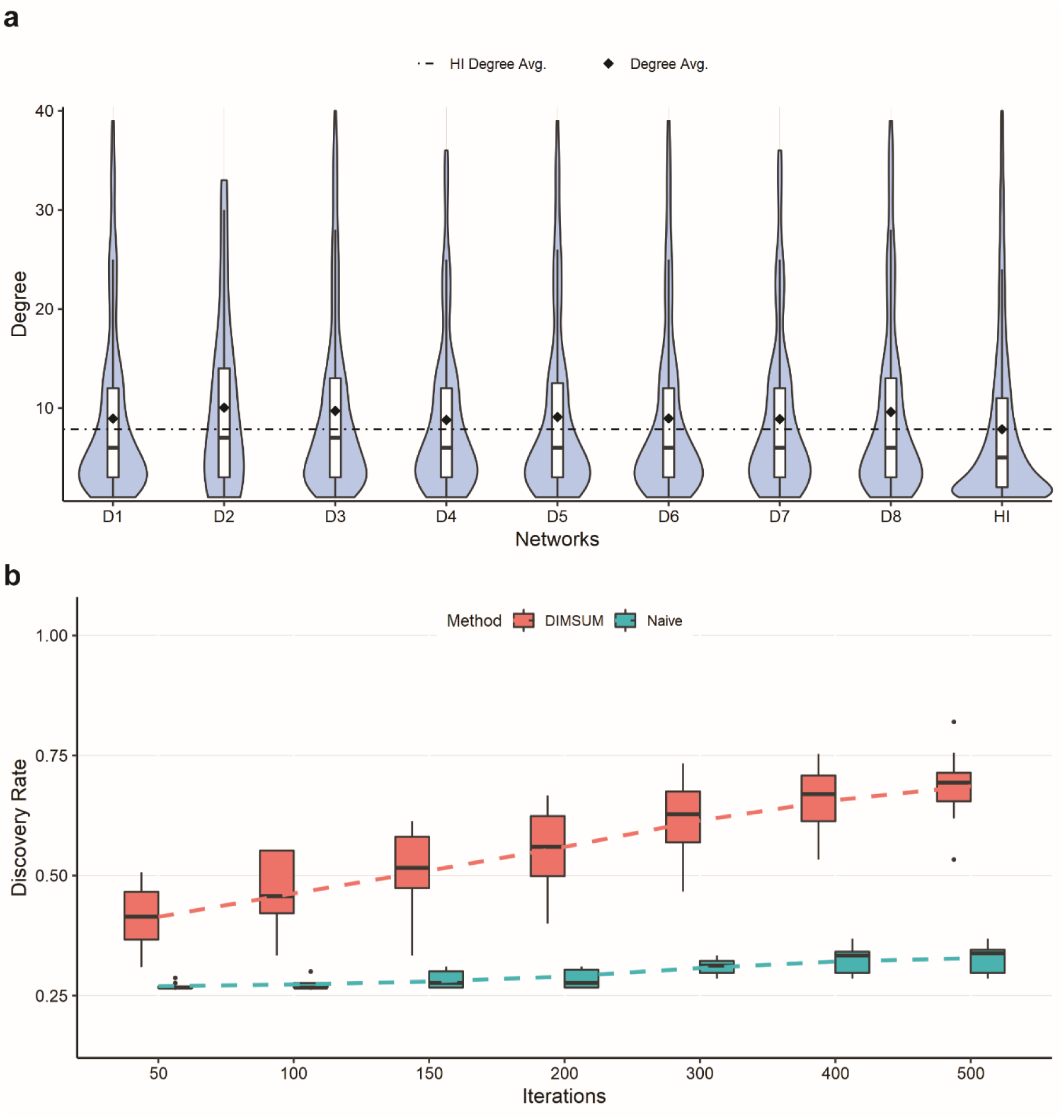
Annotated networks and comparison of DIMSUM against a naïve network propagation procedure. **(a)** The first eight violin plots represent the node degree distributions of disruptively mutated genes for eight complex diseases; the last violin plot is the node degree distribution of all genes in the human interactome (HI); the avg. degree of the disrupted genes is much greater than the avg. degree of HI, showing that highly connected genes are disrupted. **(b)** Comparison of the discovery rates of DIMSUM against a naïve network propagation procedure when randomly selecting 25% of the seed gene pool as seeds. Each box plot represents the discovery rate over all eight diseases from DIMSUM and a naïve network propagation at different iterations. DIMSUM performs significantly better than Naïve at the initial 50 iterations and improves drastically with increasing iterations.

Following annotation of the interactome with GWAS and functional data, we adopted a network propagation strategy to implicate protein interactions most likely to be influenced by disruptive SNVs and proposed a novel subnetwork extraction algorithm to find the mutation-specific module with the most “impact”. To determine if our protocol benefited from integrating GWAS data and functional annotation into the interactome, we compared our protocol against a naïve network propagation solution on the basic human interactome, i.e., without integration GWAS or functional annotation data [41, 46]. We used three selection ratios, 25%, 50% and 75%, to randomly pick the number of seeds and compared the numbers of remaining seeds that could be rediscovered after applying either DIMSUM or naïve propagation. The edges were assigned with the same weight value 1 for each edge, and the nodes were assigned with weight value 1 for the seed genes and 0 for the rest. After propagation in the naïve approach, we selected genes with the highest node score to add to the module (Fig. 2.b and Supplementary Fig. 2). The edges were assigned with the same weight value 1 for each edge, and the nodes were assigned with weight value 1 for the seed genes and 0 for the rest.

After propagation in the naïve approach, we selected genes with the highest node score to add to the module (Fig. 2.b and Supplementary Fig. 2). The results demonstrated that our protocol had a substantially higher discovery rate compared to the naïve network propagation strategy. As an additional experiment, we used the entire set of seeds for the network propagation in both methods, and then examined the top 100 discovered genes. When checking the overlap between these two gene sets, we found that the set of overlapping genes consisted of 6 genes on average across eight diseases. The results suggested that our method emphasized the genes with the greatest functional impact on the interactome and were not driven exclusively by the information propagated from the seeds. In other words, the genes extracted in the last step of our method indicated strong association with the disease and also reflected the severe damage caused by the mutations.

### 3.4. Comparison against DIAMOnD and SCA

To validate the performance of our methods, we compared our method against two seeds-based module detection methods, DIAMOnD and SCA {refs}. DIAMOnD (<u>DI</u>se<u>A</u>se <u>MO</u>dule <u>D</u>etection) is one of the most popular methods for module detections. It was developed based on the observation that the connectivity significance is a more predictive quantity characterizing the module’s interaction patterns, rather than connection density. The core idea of the algorithm was that, given a set of disease genes as seeds, it ranked all the candidates connected to the seeds based on their connectivity significance and added them to the existing seed set. SCA (Seed Connector Algorithm) is a recently developed seeds-based module detection method. SCA was built on the idea of seed connectors, which served as “bridges” of different network branches that were induced by seed genes. It selected a gene that maximally increased the size of the largest connected component of the subnetworks as the seed connector, and added them to the existing module.

We compiled a list of known disease-related genes for eight GWAS datasets as our benchmark (Supplementary Table 4). We then manually checked which of the added genes were supported with the literature evidence. We found that the lists of seed genes across eight diseases had between 5 and 29 of the disease genes supported by literature (Table 1). We next checked whether DIMSUM outperformed the other two methods in terms of the prediction accuracy in discovering the genes for our concurrence of the added gene list and the disease gene list from supported by literature. We observed that our method outperformed both DIAMOnD and SCA in all eight cases but one (osteoporosis), where DIMSUM did not find a match while both DIAMOnD and SCA found a single gene match. In fact, the predictions from DIAMOnD and SCA barely found any literature-supported disease genes (Table 1). The results also showed that except for the genes with very strong statistical signals from GWAS studies, implicating other disease genes through a network-based approach is a challenging task.

We next carried out GO enrichment analysis on each gene set using all three categories of GO terms at the third level. GO annotation was then used to find how many genes from the newly obtained module shared the same GO terms with the seed genes. The results (Fig. 3.a) showed that DIMSUM on average yielded a higher number of GO terms compared to both DIAMOnD and SCA. O the other hand, our method did not always have the highest number of shared GO terms for an individual disease. Interestingly, our analysis also showed that the dominant GO terms for genes extracted by all three methods fell in the Biological Process category.

**Figure 3.**
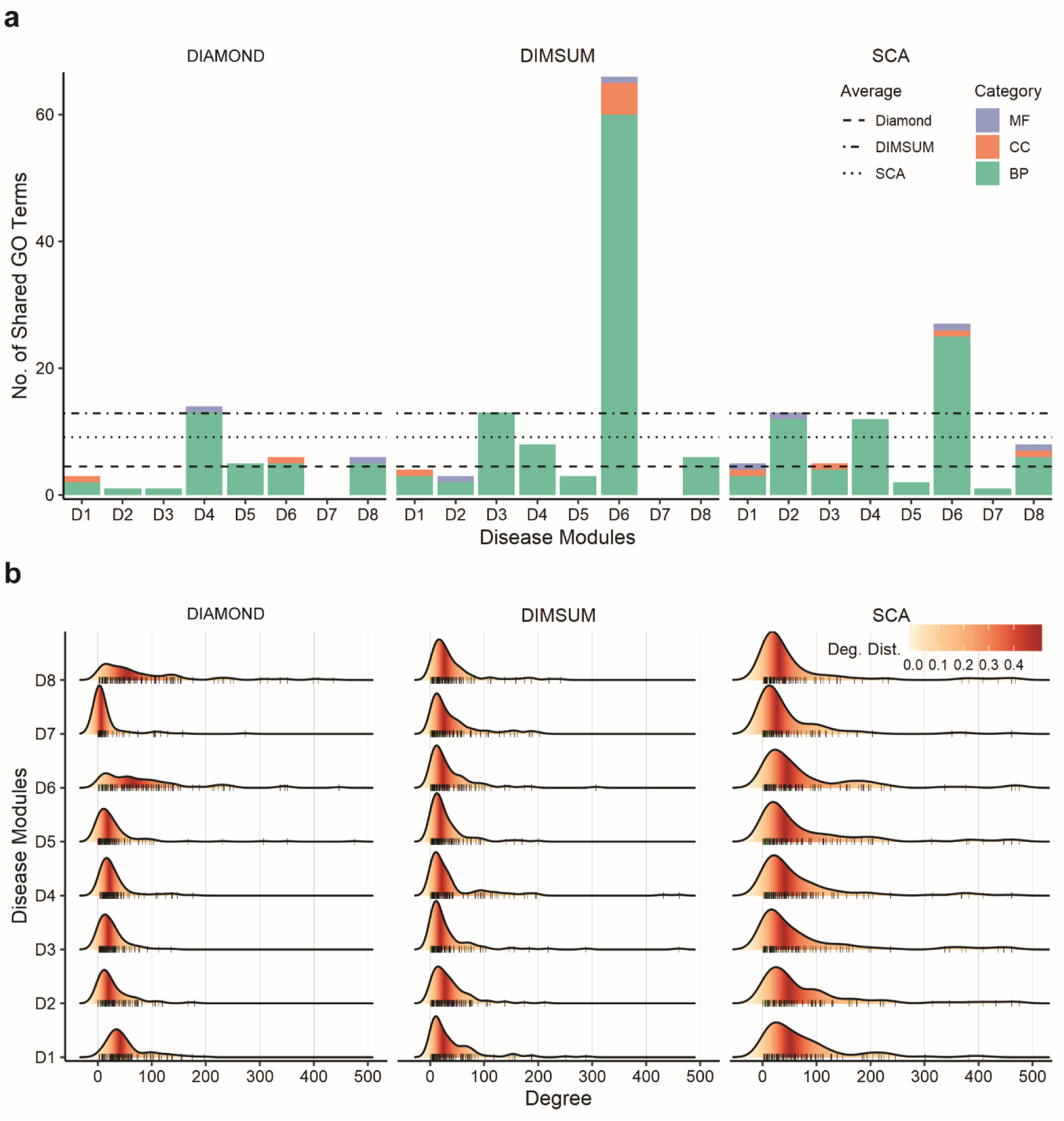
Comparison of biological relevance and topological properties of disease modules detected using three methods. (**a**) Bar plots of the number of the GO terms enriched in the added genes overlapped with the seed genes for DIAMOnD, SCA and DIMSUM for the eight disease modules. The three GO term categories are Cellular Component (CC), Molecular Function (MF) and Biological Process (BP). The average of significant and identical GO terms from DIMSUM is higher than DIAMOnD and SCA. (**b**) Degree distribution of the added genes during module detection for the eight complex diseases by each method. DIMSUM avoid bias towards high degree nodes when building the module and has lower degree for all eight modules.

Next, we examined the topological properties of the modules generated from the three methods. To quantify the structural difference of the modules generated from three methods, we focused on two topological properties. First, we calculated the connection density of the disease modules. Previous studies [25] showed that the connection density was not the primary quantity to characterize the connection patterns among disease proteins. It was further argued that, in biological networks, the paths through low-degree nodes bore stronger indications of functional similarity than the paths that went through the high-degree nodes, or hubs [47]. These findings suggested a good strategy for a module detection method should reduce the density of the detected modules and mitigate the influence of the hubs in the human interactome. When comparing these three methods, we found that DIAMOnD had the highest connection density in the detected networks, whereas modules generated from SCA and DIMSUM had much lower density (Fig. 4 and Suppl. Fig 3). We also observed that SCA favored the genes with extremely high degrees, as indicated by the genes in the long tails (Fig. 3b). The low density of the modules and node degree distributions from DIMSUM suggested that our method was not biased towards detecting interaction hubs, and thus was expected to extract genes with similar functions from the rest of the interactome more efficiently than DIAMOnD.

**Figure 4.**
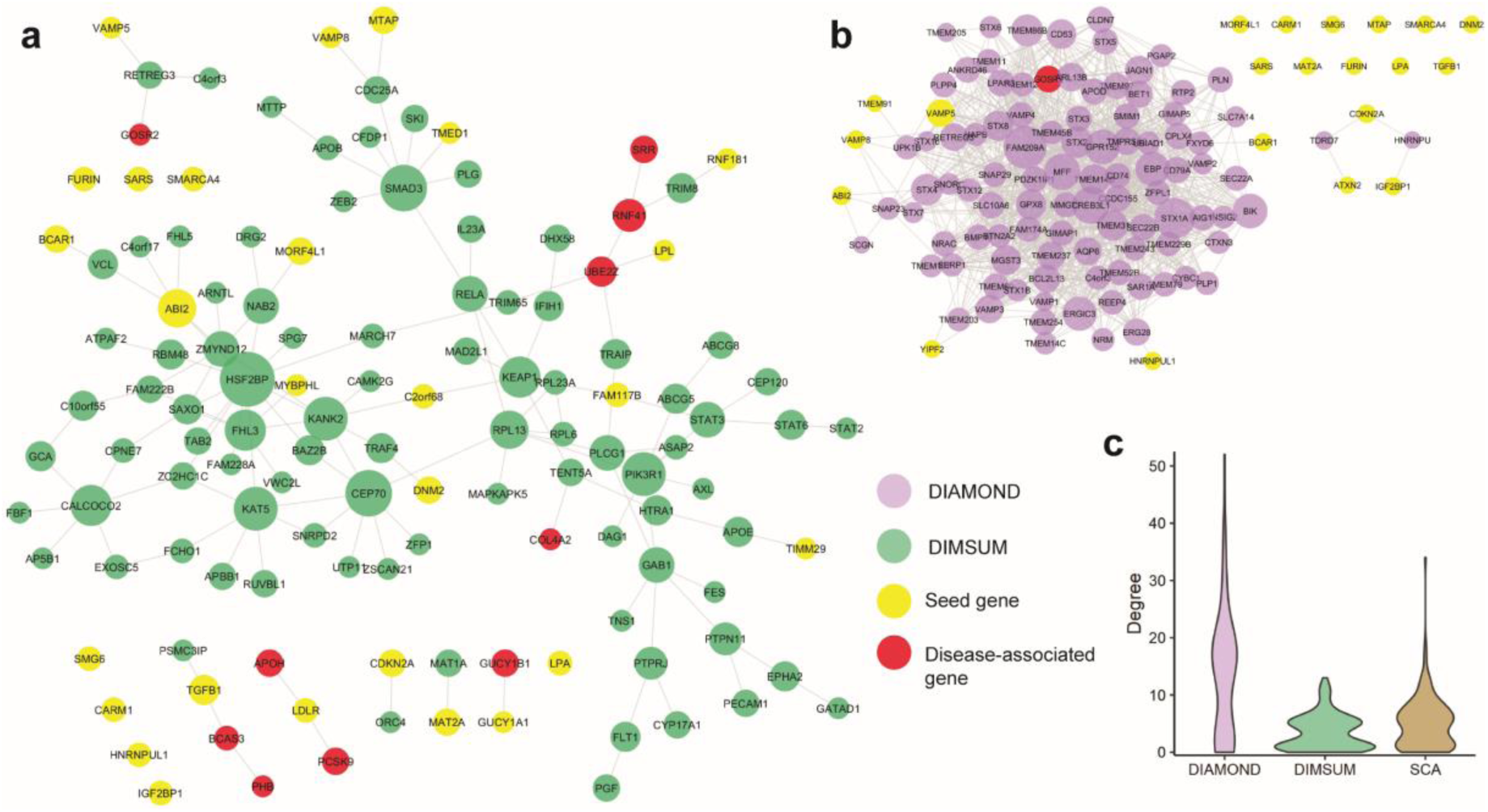
Comparison of Coronary Artery Disease modules discovered by the three methods. (**a**) Largest connected component and satellite components detected by DIMSUM, the yellow nodes represent seed genes, and the red nodes represent disease-associated gene supported by literature. (**b**) Large and high-density module detected by DIAMOnD. DIMSUM identifies ten CAD associated genes, whereas DIAMOnD identifies only one. (c) Degree distribution of the modules generated by each method, shows DIMSUM does not tend to grow a highly dense clique and it is not biased toward the hubs with very high degree.

Another interesting distinction between the methods was the fact that DIAMOnD and SCA had a tendency to grow a single giant globular component, whereas our method typically built a major component accompanied with a set of smaller, “satellite”, components. In particular, we found some of the disease-associated seed genes occurring in the small satellite subnetworks from the modules obtained by DIMSUM, but not DIAMOnD or SCA. This observation suggests that the smaller subnetworks play equally important role in defining the disease phenotype.

### 3.5. Case study 1: Coronary artery disease

As a first case study, we considered an application of DIMSUM to extract a network module centered around coronary artery disease (CAD) and compared the module to those ones derived by SCA and DIAMOnD using the same set of seed genes (Fig.4, Suppl. Fig 3). The CAD GWAS dataset was obtained from the CARDIoGRAMplusC4D Consortium[39]. After pre-processing and applying the Pascal tool, we were able to curate a seed gene pool consisting of 37 genes, 24 of which could be mapped to the human interactome. The seed genes were spread across the entire interactome, with very few direct interactions between each other. For each of the three methods we extracted 100 genes in addition to the original seed gene set to form a functional module (Fig.4 and Supplementary Fig. 3). We first validated the obtained new genes from each of the three modules against known CAD-associated genes that were collected from literature. DIMSUM outperformed both DIAMOnD and SCA (Table 1): DIAMOnD had only one and SCA reported no known CAD-related genes.

The CAD disease module from DIAMOnD formed a clique-like structure (Fig. 4b). There were dense connections inside the largest connected component, the property typically not observed in a functional module [25]. Besides, the largest component originated from only seven seed genes. During the later stage of the DIAMOnD algorithm, extraction of additional genes to form the disease module was determined by several genes, including *FAM209A, STX1A*, and *CREB3L1*, which were not seeds and which were added in the early steps of the method run, rather than from the initial seed gene pool. We surveyed the literature and did not find a strong link between these non-seed genes and CAD. We also observed that the rest of the seed genes became isolated and separated from the largest component. On the contrary, SCA generated a globular structure for the disease module, in which the largest connected component (LCC) includes most of the seed genes. This is not surprising, as SCA is specifically designed to add “seeds connector” to grow the LCC maximally, rather than the genes with functional importance. The addition of seeds connector, therefore, was biased towards the hubs in the interactome. SCA tended to add many genes with high values of node degrees, as indicated by the long tails of the degree distributions. This phenomenon was not only demonstrated in the case of CAD module, but was also evident for the other diseases we studied (Fig. 3b). However, recent work revealed that proteins connected to the high-degree hubs were less likely to have similar functions, compared to the proteins that interacted with a protein of significantly lower node degree [48].

Finally, when examining the disease module generated from DIMSUM we found that DIMSUM module included most of the seed CAD-related genes (Table 1 and Fig. 4). In addition, there was a core component in the module set which was topologically different compared to the core components generated by DIAMOnD and SCA: it did not form a highly dense clique and it was not biased toward the hubs with very high node degrees (Fig. 4.d). In addition to the core component, the DIMSUM functional module included small satellite subnetworks that harbored several functionally important genes known to be associated with CAD but not reported in the seed gene pool. Three of these satellite modules contained five genes associated with CAD, namely: *PHB, BCAS3, GOSR2, APOH* and *PCSK9*.

### 3.6. Case study 2: Schizophrenia and Bipolar disorder

In the second case study, we use DIMSUM to find if two psychiatric disorders, schizophrenia and bipolar disorder, that shared symptoms also shared functional modules. Bipolar disorder (BPD), is a mental disorder, also known as manic-depressive illness, that causes unusual shifts in mood, energy and activity levels, often resulting in periods of depression or mania [49, 50]. Schizophrenia (SCZ) is a chronic and severe mental disorder that is represented by abnormal behavior and an altered notion of reality where the patients hear voices or see objects/persons that are not real [51]. While schizophrenia is not as common as other mental disorders, the symptoms can be very disabling. Schizophrenia and bipolar disorder had many common traits previously documented [52, 53].

The DIMSUM algorithm was supplied with 76 seed genes for Schizophrenia and 15 seed genes with Bipolar Disorder extracted and processed from two GWAS studies [54, 55]. There was no overlap between the seed gene sets for the two diseases. For each disease, additional 100 genes were extracted by DIMSUM to form the disease-centered module. We first queried the genes from the obtained BPD module against a list of BPD-associated genes from another recently published GWAS of the Psychiatric Genomic Consortium Bipolar Disorder Working Group [56]. As a result, we identified four genes from the module that were not among the initial set of seed genes but were found in the above GWAS study by Bipolar Disorder Working Group: *RIMS1, ERBB2, STK4* and *MAD1L1*. These four genes have been previously shown to play functional roles in a number of neurological disorders [57-59]. For example, *RIMS1* is a *RAS* superfamily member, and the encoded protein regulates synaptic vesicle exocytosis [60]. Mutations occurring on RIMS1 genes have been suggested to play a central role in cognition[61]. In addition to BPD, it is associated with autism spectrum disorder, neurodevelopmental disorders, and intellectual disability [57, 62].

To determine disease-associated genes in the SCZ module we relied on a recent study that categorized the disease associated genes under three tiers based on diagnosis, polygenic risk scores (PRS) and those reported by the Psychiatric Genomic Consortium (PGC) [63]. In total, we found that 33 genes among the 100 added genes were present in the PGC gene set, of which four were in Tier 1 (*CDK2AP1, MDK, ZFYVE21* and *RRAS*). Genes in this Tier were found to be significantly associated with both diagnosis and PRS. Perhaps the most interesting of these four was MDK, a gene associated with many important neurological processes, *e.g*., cerebral cortex development, behavioral fear response, short-term memory, and regulation of behavior [64, 65]. Eight more genes belonged to Tier 2, *i.e*., associated with diagnosis but not PRS.

Finally, we determined if the BPD and SCZ modules shared any genes—or more importantly submodules—in common. It had been established that bipolar disorder and schizophrenia shared a large overlap of genetic risk loci and often exhibited similar symptoms like mania and depression [53]. Thus, in spite of the missing overlap between the two seed genes sets between those disease, the modules enriched with more disease-related genes could share common genes. Intriguingly, the BPD and SCZ modules were found to share a smaller sub-module of 10 genes connected with each other and with other BPD and SCZ genes (Fig. 5a). Out of 10 genes found shared between BPD and SCZ modules, four genes (*HIST1H3A, PBX4, MAD1L1, NRAS*) were known to be strongly associated to both disorders (Fig 5a). We conjectured that the common genes between the BPD and SCZ modules could provide insights into the phenotypic similarities between the two diseases. To support this hypothesis, we revisited the functional predictions from SNP-IN tool involving these ten genes. We found that while the mutations occurring in both diseases were quite diverse, a small group of mutations targeted the same subnetwork centering around HIST1H3A and HIST1H4A (Fig. 5b). Both of these genes were the core units of the nucleosome, implicated in a number of neuro-psychiatric disorders [66, 67]. Most of the mutations occurring in this subnetwork were predicted by SNP-IN tool to disrupt the corresponding PPIs (Fig 5c, Supplementary Table 5). Thus, different mutation frequencies and combinations for BPD and SCZ could give rise to different rewiring, or edgetic, effects of the HIST1H3A/HIST1H4A centric subnetwork. These results lead us to suggest that the different rewiring patterns of the same subnetwork could explain the phenotypic similarity but underlie differences in symptomatic severity between the two diseases.

**Figure 5.**
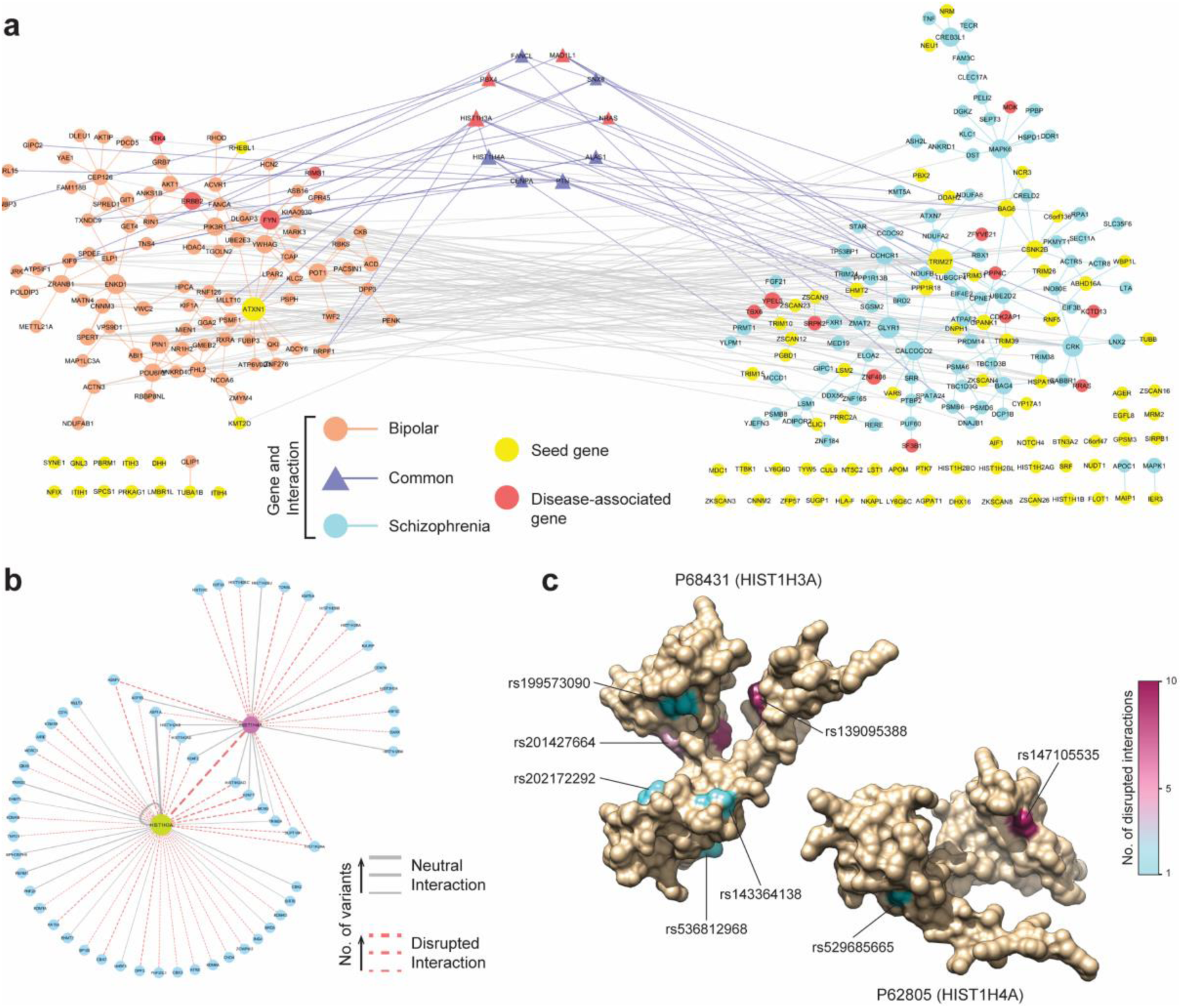
Analysis of Bipolar disorder and Schizophrenia modules discovered by DIMSUM. **(a)** The BPD and SCZ modules are represented by the left and right components respectively, and the genes common to both disorders are in the small central component. DIMSUM discovers a total of nineteen disease-associated genes in both modules. (b) The rewiring of the subnetwork centered around the shared histone genes HIST1H3A and HIST1H4A. The dash line indicates the accumulative damage based on the SNP-IN predictions. (c) Protein structures for the histone genes HIST1H3A and HIST1H4A on the left and right respectively. Disruptive mutations occurring on these two genes are observed in both BPD and SCZ; HIST1H3A carries six disruptive mutations and HIST1H4A carries two. The color of the mutated residue corresponds to the number of interactions it disrupts.

## 4. Discussion

In this work, we proposed a computational framework for functional disease module detection, DIMSUM, which integrates GWAS datasets with the human interactome, propagating the functional impact of nsSNVs. Our module detection approach first annotates the network with the functional information from the genes and associated mutations followed by the network propagation to determine new genes associated with the same disease and, finally, subnetwork extraction. We assessed our approach using a set of 8 complex diseases against two state-of-the-art seed-based module detection methods, DIAMOnD and SCA. The integration of multiple data types within a single analysis framework allowed us to improve the effectiveness of module detection. In particular, the evaluation results showed that DIMSUM outperformed both DIAMOnD and SCA—our approach was able to yield modules with stronger disease association and more biological relevance.

Integrating biomolecular networks across various types of data, including -omics profiles, GWAS, and functional annotation, have proven to be powerful for the detection and interpretation of biological modules [29]. GWAS investigates the entire genome and identifies the genomic loci related to a disease, providing a “macro view” of the underlying genetic architecture. On the other hand, leveraging functional annotation tools like SNP-IN tool, enables a “micro view” by examining the specific and localized mechanistic effect of mutations providing insights into disease etiology. Our analysis framework facilitates joint interpretation of the biological information originating from those two different perspectives. Furthermore, the network propagation procedure allows to interpret the list of candidate genes into a genome-wide spectrum of gene scores reflecting the disease association signal. The ability to amplify the signals from the seed gene pool by our approach has been previously proven to be helpful when identifying the genetic modules that underlie human diseases [20].

The case studies of eight complex diseases showed that the final modules obtained by DIMSUM typically include a large connected component containing most of the genes associated with a disease. This capacity of the algorithm to merge the initially isolated seed genes into a connected core component may prove to be helpful for understanding of molecular mechanisms underlying the complex disease that are often carried out by functioning of molecular complexes and pathways, rather than isolated proteins. Furthermore, the submodules disconnected from the largest component were also found to harbor a considerable number of genes related to the disease. Examination of the discovered modules and findings from the case studies prompted us with a new hypothesis that underlie the importance of the system-wide variation effects for complex genetic diseases. Specifically, we hypothesize that disease phenotypes observed in complex diseases, such as coronary artery disease and schizophrenia, may be a consequence of rewiring of an orchestrated functional module system rather than the abnormal functioning of the independent genes. Such functional module system would consist of a core module and several smaller satellite modules. The core unit is mainly responsible for the disease, while rewiring of the smaller satellite modules could also contribute to the disease progress and disease phenotype diversity. We further hypothesize that some satellite modules could correspond to a specific symptom node in the recently proposed symptom network for the psychiatric disorders [68]. Thus, the specific rewiring of the functional module system could help explaining the disease subtypes or different symptom combinations among patients.

## Supplementary Materials

The following are available online. Figure 1: Illustration of methodology for structure-based prediction of SNP’s effect on PPI when applying SNP-IN tool; Figure 2: Comparisons of the discovery rates of our approach against a naïve network propagation procedure when randomly selecting 50% and 75% from the seed gene pool as seeds; Figure 3: Coronary artery disease (CAD) module discovered by the SCA algorithm; Table 1: Description of the eight GWAS datasets curated for this work; Table 2: Seeds generated from Pascal tools from GWAS datasets for the eight complex diseases; Table 3: SNP-IN tool annotation results for the eight GWAS datasets; Table 4: Disease-gene association data curated from OMIM, HGMD; Table 5: HIST1H4A-HIST1H3A centered PPI subnetwork and associated disruptive mutations.

## Author Contributions

H.C. conceived the idea. H.C., S.S. and D.K. designed the experiment(s), H.C. and S.S. conducted the experiment(s). H.C., S.S. and D.K. analyzed the results, and H.C., S.S. and D.K. wrote the manuscript. All authors reviewed the manuscript.

## Funding

“This research was funded by National Science Foundation (1458267) and National Institute of Health (LM012772-01A1) to D.K.”

## Conflicts of Interest

The authors declare no conflict of interest.

