## Supplementary materials and figures for "Enriching human interactome with functional mutations to detect high-impact network modules underlying complex diseases"

### Supplementary Information

#### List of figures:


| | |
|---|---|
| **Supplementary Figure 1** | Structure-based prediction of SNP's effect on PPI when applying SNP-IN tool. |
| **Supplementary Figure 2** | Comparisons of the discovery rates of our approach against a naïve network propagation approach. |
| **Supplementary Figure 3** | Coronary artery disease (CAD) module discovered by the SCA algorithm. |


#### List of tables:


| | |
|---|---|
| **Supplementary Table 1** | Description of 8 GWAS datasets applied in this study. |
| **Supplementary Table 2** | Seeds generated by the Pascal tool from 8 GWAS datasets of complex diseases. |
| **Supplementary Table 3** | SNP-IN tool annotation results for 8 GWAS datasets. |
| **Supplementary Table 4** | Disease gene association data curated from OMIM, HGMD and GeneCards. |
| **Supplementary Table 5** | HIST1H4A-HIST1H3A centered PPI subnetwork and associated disruptive mutations |

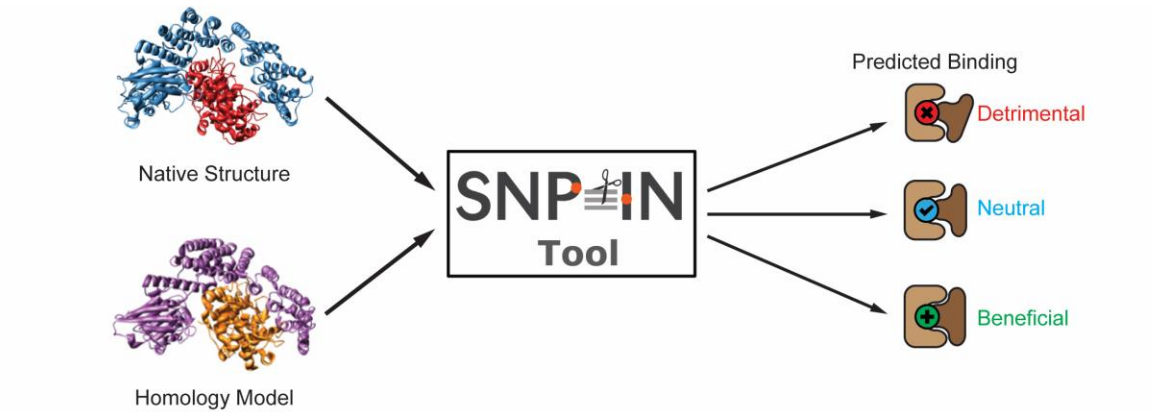


**Supplementary Figure 1. Structure-based prediction of SNP's effect on PPI when applying SNP-IN tool.** The SNP-IN tool uses native structure (when available) or a homology model of the PPI as an input. The output is the predicted PPI-rewiring effect of the SNP with three possible outcomes: (i) Detrimental: where the binding is lost; (ii) Neutral: the binding is preserved and (iii) Beneficial: the binding is strengthened.

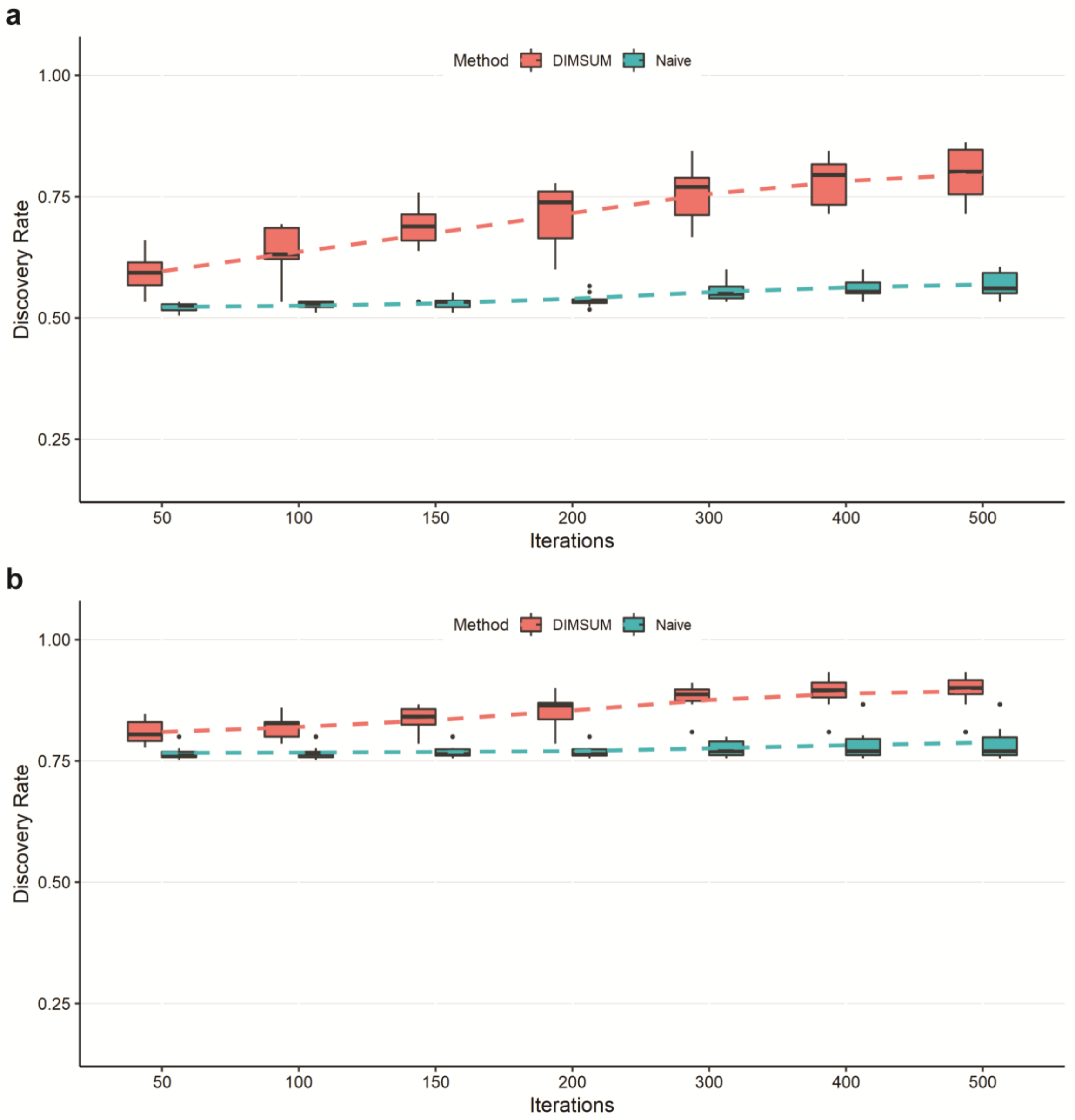


**Supplementary Figure 2. Comparisons of the discovery rates between our approach and a naïve network propagation approach.** (**a**) 50% of nodes are randomly selected from the seed gene pool where DIMSUM has a greater trend in discovery rate with respect to the increasing number of iterations reaching an average of 0.75. (**b**) 75% of nodes are randomly selected from the seed gene pool where DIMSUM again outperforms the Naïve method, reaching an average discovery rate of 0.9 with 500 iterations.

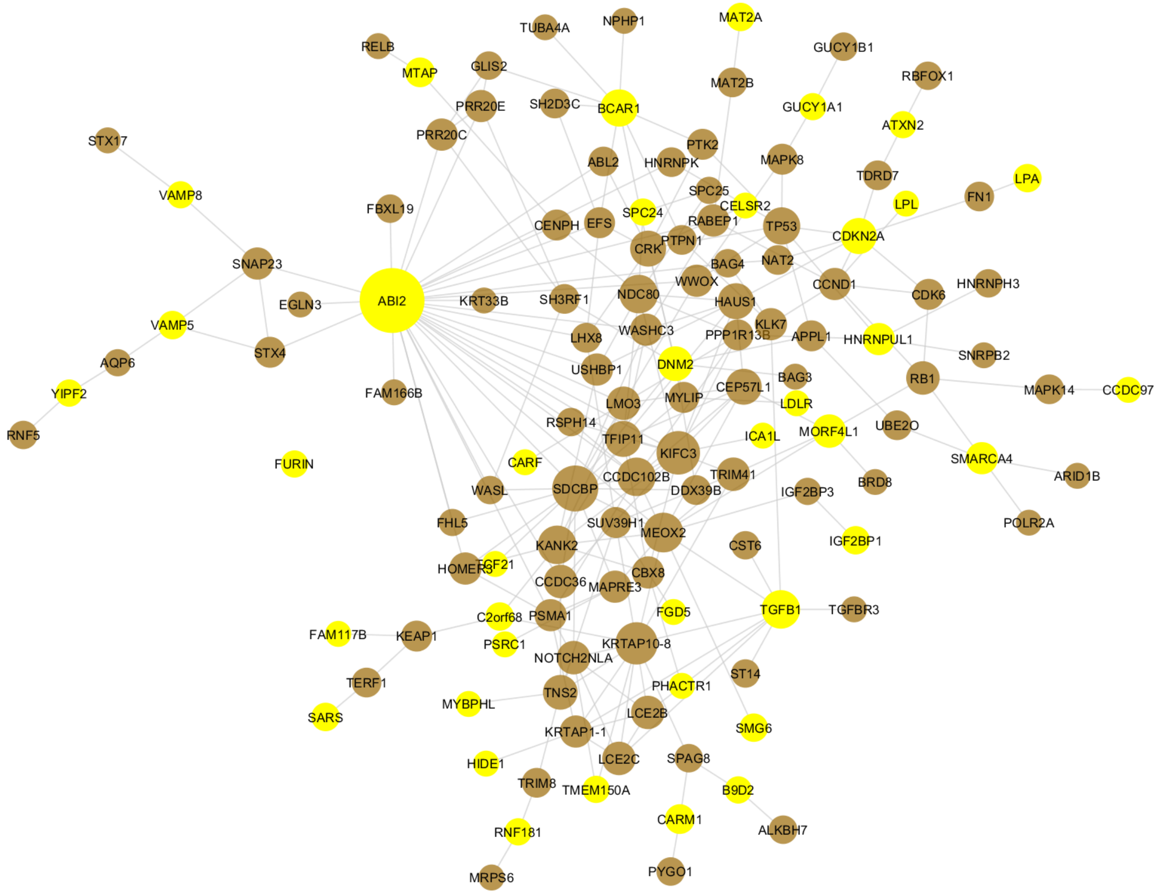


**Supplementary Figure 3. Coronary artery disease (CAD) module discovered by the SCA algorithm.** Yellow nodes represent the seed genes for CAD and brown nodes represent the added genes. The SCA algorithm tries to include as many seeds as possible when building the module. No CAD associated genes were discovered by SCA, and the degree distribution was greater (i.e. high degree nodes or hubs were added) than in the module discovered by DIMSUM (Fig. 4).
